# Image Quality Causes Substantial Bias in Three-Dimensional Speckle-Tracking Echocardiography Measures

**DOI:** 10.1101/500777

**Authors:** Lamia Al Saikhan, Chloe Park, Alun Hughes

**Affiliations:** Institute of Cardiovascular Science, School of Life and Medical Sciences, University College London, London, United Kingdom; Department of Cardiac Technology, College of Applied Medial Sciences, Imam Abdulrahman Bin Faisal University, Dammam, Kingdom of Saudi Arabia; MRC Unit for Lifelong Health and Ageing, University College London, London, United Kingdom.

**Author notes:** **Correspondence**: Lamia Khaled Al Saikhan, Institute of Cardiovascular Science, University College London, Gower Street, London WC1E 6BT, Twitter: @Lamya_k_s, Tweet: Sub-optimal 3DE image quality introduces a substantial systematic bias on LV deformation and rotational indices by 3D-STE.

**Keywords:** 3D speckle tracking, sub-optimal image quality, strain, rotation

## Abstract

**BACKGROUND:** Three-dimensional speckle-tracking echocardiography (3D-STE) is increasingly used to assess left ventricular (LV) mechanics but the quantitative effect of image quality on measurements is not known.

**OBJECTIVES:** To assess the impact of image quality on 3D-STE derived LV indices.

**METHODS:** Echocardiography was performed in two groups of 18 healthy participants. In the first study, optimal and intentionally poor-quality images were acquired. In the second study, a sheet of ultrasound-attenuating material (neoprene rubber) of three different thicknesses (2, 3 and 4 mm) was used to mimic mildly, moderately and severely impaired image quality respectively.

**RESULTS:** In both studies sub-optimal image quality resulted in a systematic underestimation bias in all LV deformation and rotational indices. LV ejection fraction and volumes were also consistently underestimated. The extent of the bias was proportional to the impairment in image quality (i.e. the poorer the image quality the larger the bias). Reproducibility was also less good for sub-optimal images, although LV volumes and ejection fraction showed excellent reproducibility irrespective of image quality.

**CONCLUSIONS:** Sub-optimal image quality introduces a substantial systematic bias and impairs the reproducibility of 3D-STE. Bias related to image quality might have important clinical implications since its magnitude is similar to that reported in association with disease and may confound associations between disease and LV mechanics.

**CONDENSED ABSTRACT:** Three-dimensional speckle-tracking echocardiography (3D-STE) is increasingly used in clinical practice. We assessed the impact of sub-optimal image quality on left ventricular (LV) deformation indices either by poor echocardiographic technique or by impairing ultrasound propagation using a neoprene sheet. Sub-optimal image quality introduces a substantial systematic bias that increases with poorer image quality. Bias due to image quality might have important clinical implications since its magnitude is similar to that reported in association with disease.

## INTRODUCTION

Accurate assessment of left ventricular (LV) function is clinically important to the determination of prognosis and therapeutic strategies (1,2). Currently, assessment is mainly based on global volume-based measures and the most commonly used measure is LV ejection fraction (LVEF) measured by echocardiography (2,3). Despite reductions in LVEF being associated with poor prognosis (4–9), LVEF has a number of limitations, including load dependency, poor reproducibility, and being more an indicator of circumferential/radial function than longitudinal function (2,3). It is well recognised that impaired LV systolic function may occur when the LVEF is still normal (3).

Myocardial deformation imaging has emerged as a promising tool to quantify myocardial mechanics. It provides detailed characterization of LV performance, namely deformation in 3-dimensions (longitudinal, circumferential and radial), in addition to providing information on torsion and twist (2,10). While tissue-Doppler imaging was first used to assess myocardial deformation, this method is angle dependant and has been largely superseded by speckle-tracking echocardiography (STE) (10). This technique is based on tracking myocardial features of a grey scale B-mode image called *‘speckles’*, which are generated by the interaction between the ultrasound beam and the myocardium, providing information about the global and regional myocardial function (10–12). STE can be performed using twodimensional STE (2D-STE), but this has the limitation that tracked speckles are assumed not to move ‘out of plane’ during LV motion (10,13,14). Three-dimensional STE (3D-STE) has recently emerged and may overcome some of the limitations of 2D-STE. Theoretically, 3D-STE should be more accurate as blocks of speckles can be tracked irrespective of their direction, overcoming the ‘out of plane’ motion phenomenon (10,13,14). 3D-STE also allows LV myocardial deformation to be assessed simultaneously in all directions, overcoming the heart rate variability problem associated with the multiple acquisitions required for 2D-STE (10,13–15). For these reasons, in theory, LV twist can also be calculated more precisely with 3D-STE than with 2D-STE (15,16); however, the comparatively low spatial and temporal resolutions of 3D-STE are currently major concerns (17–19).

Sicker/frailer patients often have sub-optimal echocardiographic image quality, which may limit the utility of 3D-STE. While it is widely believed that 3D-STE derived LV deformation indices are influenced by image quality, quantitative evidence on this is limited. Previous studies have evaluated the impact of image quality by excluding individuals with sub-optimal images and examining the agreement between 3D-STE and other methods such as 2D-STE (20) or inter-vendor agreement (21). Alternatively, within study associations between 3D-STE derived LV deformation indices and image quality have been reported (22), but the interpretation of these observations is difficult due to potential confounding resulting from the associations between aging, obesity and disease with image quality. As yet, no study has provided quantitative evidence of the unconfounded impact of sub-optimal image quality on 3D-STE derived LV indices. Therefore, we performed a study to quantitate the impact of intentionally distorted image quality on LV deformation indices by 3D-SET, with the aim of 1) establishing whether sub-optimal image quality causes a systematic bias; 2) quantifying the extent of bias in proportion to the impairment in image quality; and 3) determining the effect of sub-optimal image quality on within participant reliability of 3D-STE derived LV indices.

## METHODS

### STUDY POPULATION

This study consists of two sub-studies:

1. Healthy participants (n=23) with no previous cardiac medical history were screened prior to undergoing three-dimensional echocardiography (3DE) to examine the effect of intentionally poor image acquisition technique on LV deformation indices. Participants with sub-optimal echocardiographic window on scanning were excluded (n=5), leaving a final sample size of 18.
2. An additional group of healthy participants (n=21) was recruited to quantify the impact of degrading image quality on LV deformation indices by impairing ultrasound propagation using an attenuating material (analogous to an unfavourable body habitus). Three screened participants had poor echocardiographic windows and were excluded leaving a final sample size of 18.

The protocol was approved by the institutional review board and informed consent was obtained from all participants prior to examination.

### IMAGING

Imaging was performed by a single cardiac sonographer using an EPIQ 7 ultrasound machine (Philips Medical Systems, Andover, MA) equipped with Xmatrix-array transducer (X5-1). Participants were scanned in left lateral decubitus position using a standard protocol (23). Harmonic imaging and multiple-beat 3DE mode were used. In all participants, 4 wedged-shaped sub volumes were acquired over 4 consecutive cardiac cycles during a single breath-hold. During the acquisition, special care was taken to include the entire LV cavity within the pyramidal sector volume.

In study 1, two gated wide angled 3DE full-volume datasets were obtained per participant from the apical window. The acquisition of the first 3DE full-volume dataset was performed according to EAE/ASE recommendations (23). Machine settings, including gain, sector width, and depth were adjusted by the operator to maximize the quality of images ensuring clear visualization of LV endocardial borders and avoiding echo drop out. A good 3DE image was defined as clear visualization of the endocardium in all 16 segments in both end-diastolic and end-systolic frames. The acquisition of the second dataset, on the other hand, was captured after intentionally impairing the image quality with sub-optimal echo technique. Quality was impaired by including images with echo drop out, shadow artefacts, or poor visualization of the endocardium in multiple segments throughout the cardiac cycle (Figure 1.A). Sub-optimal 3DE image was defined as the presence of at least one of the following: (1) poor visualization of the endocardium in multiple segments throughout the cardiac cycle in a 16-segment LV model, (2) the presence of echo dropout, and (3) shadow artefacts. The acquisition protocol was repeated on the same day following the same approach to assess the test-retest repeatability.

**FIGURE 1.**
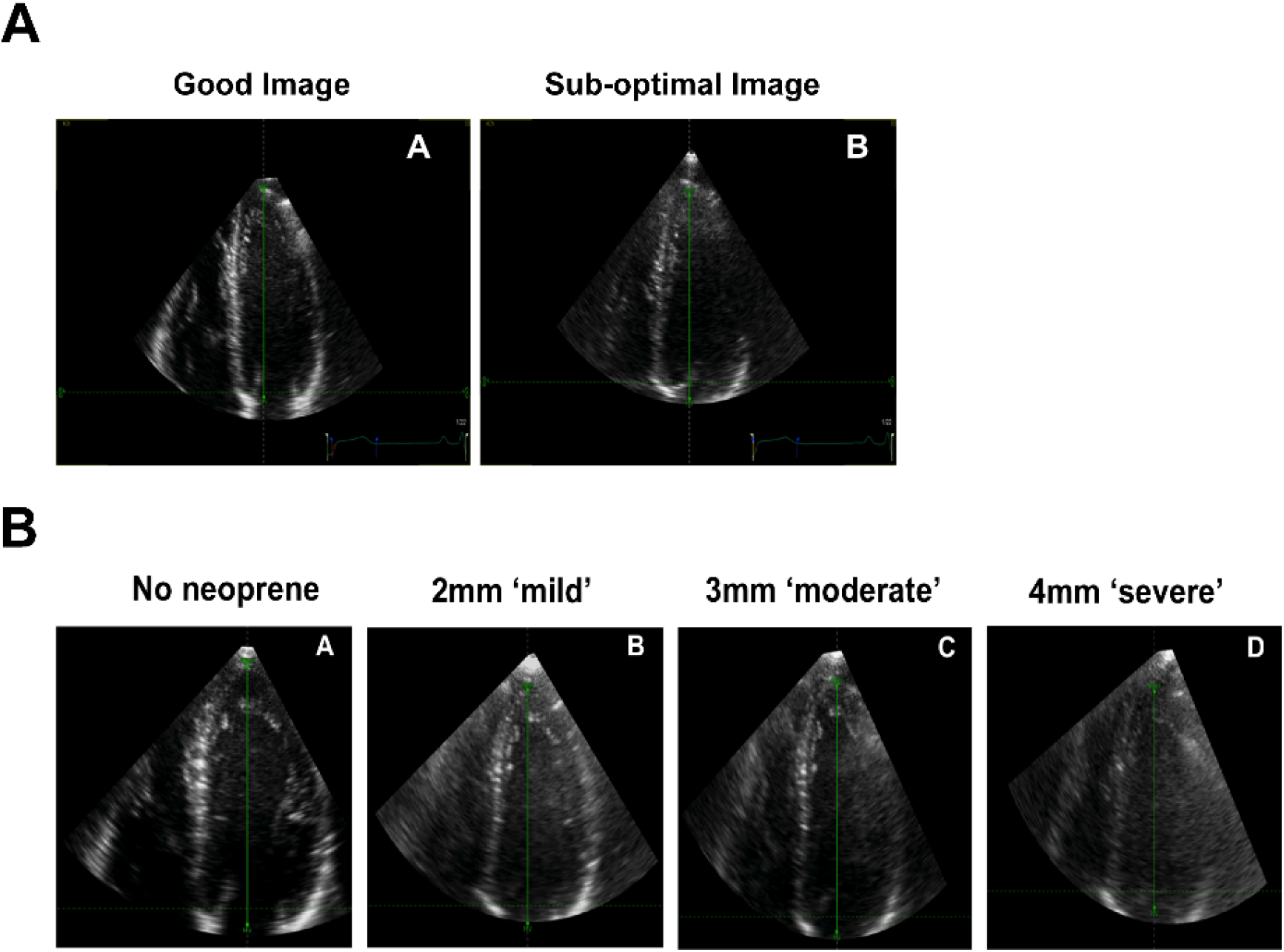
Examples of impaired 3D echocardiographic (3DE) image quality. An example of a good and sub-optimal 3DE image quality obtained from the same participant in study 1 (**A**). An example of a 3DE with an optimal quality reference (no neoprene), mild (2mm neoprene), moderate (3mm neoprene), and severe (4mm neoprene) impairment of 3DE image quality obtained from the same participant in study 2 (**B**).

In study 2, the quality of the 3DE images was impaired in graded manner using sheets of ultrasound-attenuating material, ***neoprene rubber***, of three different thicknesses (2, 3 and 4 mm) to mimic mild, moderate and severe impairment in image quality respectively (Figure 1.B). Neoprene (a polymer of chloroprene, 2 chloro-1, and 3-butadiene) was chosen as many of its acoustic properties are similar to soft biological tissues, it is durable, and it has a comparatively high attenuation coefficient (24,25). The acquisition was performed after placing the neoprene rubber between the skin and the transducer with ultrasound gel on both sides. 4 gated 3DE full-volume datasets (an optimal quality reference and 3 progressively impaired images were acquired per participant) (Figure 1.B).

All acquisitions were free of stitching artefacts with good quality ECG signals. The frame rate was maintained constant with a minimum acquisition rate of 18 frames/second (19).

### IMAGE ANALYSIS

Images were analysed using 4D LV-Analysis© software (TomTec Imaging Systems GmbH, Germany, 2015) by a single reader blinded to condition and sequence. LV deformation analysis was performed in all datasets obtained per participant (i.e. 4 analyses/participant for both sub-studies). The software automatically selected and displayed three standard apical views and one short-axis view. Alignment of the longitudinal axis of the LV in all apical views were further modified manually if needed using two anatomical landmarks at both ends (i.e. the mitral valve annulus and the apex). The endocardial borders were then defined automatically by the software in all three apical views at end-diastole. Manual adjustments could be made but these were kept as minimal as possible. The software then tracked the endocardium throughout the cardiac cycle in 3D space from which the 3D LV endocardial shell was constructed. The tracing of LV endocardial boundaries was further adjusted manually when needed in end-diastolic and end-systolic frames in all views The software then divided the LV into 16 segments using a standard 16-segment model (26) and provided curves as well as maps of global and segmental volumetric and deformation indices.

The following LV 3D-STE indices were obtained and evaluated: (1) LVEF and volumes; (2) LV rotational indices (basal and apical rotations, twist, torsion, and twisting and untwisting rates); (3) LV global longitudinal (GLS) and circumferential (GCS) strains; and (4) peak averaged segmental longitudinal (LS), circumferential (CS), radial (RS) and principle tangential (PTS) strains (a composite LV deformation index including both LS and CS). Time derivative of twist was calculated using the curve ‘row’ data to derive twisting and untwisting rate measures. From this, minimum and maximum twist rates were calculated and considered as peak untwisting and twisting rate measures respectively.

### STATISTICAL METHODS

Continuous variables are presented as mean ± standard deviation (SD). Categorical variables are presented as counts and percentages. Mixed linear modelling was used to identify systematic differences in 3D-STE LV indices. Test-retest reliability was assessed using Intraclass correlation coefficient (ICC). Reliability was defined as follows: ICC < 0.4=poor reproducibility, 0.4 ≥ ICC < 0.75=fair to good reproducibility, and ICC ≥ 0.75=excellent reproducibility (27). The sample size was chosen to ensure a width of the 95% confidence interval for the ICC equal to 0.3. For this, a minimum of 14 participants with 4 repeated measures were required to detect a bias greater or equal to one standard deviation of the measure at alpha = 0.05 with 96% power. All analyses were performed in Stata version 15.1 (StataCorp LLC, USA).

## RESULTS

### STUDY POPULATION

Participant characteristics for both studies are summarized in Table 1. The mean age was 28 (SD 6) years and 31 (SD 6), and 10 (55.5%) and 15 (83.3%) participants were male in the first and second studies respectively. In study 1, the frame rate per cycle was 21 (SD 4) and 21 (SD) for good and sub-optimal quality images respectively. For study 2, frame rates per cycle were 21 (SD 3), 21 (SD 3), 21 (SD 3) and 21 (SD 3) for the optimal, mildly, moderately, and severely impaired quality images respectively.

**Table 1.** Study population characteristics

| **Table 1 Study population characteristics** |  |  |
| --- | --- | --- |
|  | **Study 1 (n=18)** | **Study 2 (n=18)** |
| Age, year | 28±6 | 31±6 |
| Male, n (%) | 10 (55.5%) | 15 (83.3%) |
| Systolic blood pressure, mmHg | 118.2±8.6 | 123.2±9.2 |
| Diastolic blood pressure, mmHg | 73.5±7.3 | 77.3±9.5 |
| Heart rate, bpm | 72±13.8 | 69.1±14.2 |
| Height, cm | 169.9±9.4 | 172.2±8.7 |
| Weight, kg | 70.9±16.1 | 73.0±8.1 |
| Body mass index, kg/m<sup>2</sup> | 24.5±5.8 | 24.7±3.6 |
| Body surface area, m<sup>2</sup> | 1.8±0.21 | 1.8±0.12 |

ta are mean ± SD or n (%).

### THE IMPACT OF SUB-OPTIMAL IMAGE QUALITY DUE TO INTENTIONALLY POOR TECHNQIUE (STUDY 1)

Sub-optimal image quality in study 1 resulted in a systematic bias in all global and averaged segmental LV deformation indices as well as LV rotational indices (Table 2). The extent of bias was approximately 10% for strain indices and more for twist and torsion (~20%) (Figure 2). LVEF and most of LV volumes were also consistently underestimated (Table 2). No evidence of bias was observed in twisting and untwisting rate indices. Reproducibility from test-retest was excellent for LVEF and volumes, fair to good for all global and averaged segmental LV deformation indices and poor for rotational indices irrespective of whether good and sub-optimal images were analysed separately or together, but reproducibility was improved for all indices when restricting the analysis to non-degraded images [i.e. optimal quality images] (Table 2; Figure 3).

**Table 2.** Comparison of 3D-STE LV derived indices according to the image quality (study 1)

ta are mean ± SD or n (%). Da
|  |  |  |  |  |  |  |  |
| --- | --- | --- | --- | --- | --- | --- | --- |
|  | **Mean (95% CI)** |  | **Bias** |  | **ICC** |  |  |
|  | **Good** | **Sub-optimal** | **Mean $\Delta$ (95% CI)** | **P<sub>Bon</sub>** | **Combined<sup>†</sup>** | **Good<sup>‡</sup>** | **Sub-optimal<sup>§</sup>** |
| **Global LV volumetric indices** |  |  |  |  |  |  |  |
| EDV, ml | 123.9 (115.4, 132.4) | 117.7 (109.2, 126.3) | 6.1 (3.7, 8.6) | <0.0001 | 0.91 | 0.97 | 0.91 |
| ESV, ml | 53.6 (49.0, 58.2) | 53.8 (49.2, 58.4) | -0.22 (-1.5, 1.0) | >0.9 | 0.92 | 0.96 | 0.89 |
| EF, % | 56.9 (55.6, 58.2) | 54.3 (53.0, 55.5) | 2.6 (2.0, 3.2) | <0.0001 | 0.78 | 0.94 | 0.78 |
| SV, ml | 70.2 (65.8, 74.7) | 63.8 (59.4, 68.3) | 6.4 (4.8, 7.9) | <0.0001 | 0.88 | 0.97 | 0.90 |
| **Global and average segmental\* LV deformation indices** |  |  |  |  |  |  |  |
| GCS, % | -28.0 (-29.0, -27.0) | -25.7 (-26.7, -24.7) | -2.3 (-2.9, -1.7) | <0.0001 | 0.72 | 0.88 | 0.52 |
| GLS, % | -21.4 (-22.2, -20.5) | -20.2 (-21.1, -19.3) | -1.1 (-1.8, -0.47) | 0.002 | 0.50 | 0.62 | 0.41 |
| Peak CS, % | -28.2 (-29.3, -27.2) | -25.7 (-26.7, -24.6) | -2.5 (-3.1, -1.9) | <0.0001 | 0.72 | 0.86 | 0.54 |
| Peak LS, % | -20.4 (-21.2, -19.5) | -19.6 (-20.5, -18.8) | -0.73 (-1.4, -0.02) | 0.172 | 0.47 | 0.66 | 0.39 |
| Peak PTS, % | -33.0 (-34.1, -31.9) | -31.4 (-32.5, -30.3) | -1.6 (-2.3, -0.86) | <0.0001 | 0.62 | 0.82 | 0.36 |
| Peak RS, % | 40.9 (39.6, 42.1) | 38.2 (36.9, 39.4) | 2.6 (1.98, 3.3) | <0.0001 | 0.72 | 0.80 | 0.51 |
| **LV rotational indices** |  |  |  |  |  |  |  |
| Peak basal rotation, | -8.0 (-9.7, -6.2) | -6.7 (-8.5, -4.9) | -1.2 (-2.7, 0.22) | 0.388 | 0.48 | 0.54 | 0.60 |
| Peak apical rotation | 6.2 (5.2, 7.3) | 4.6 (3.5, 5.7) | 1.6 (0.61, 2.6) | 0.008 | 0.36 | 0.41 | 0.24 |
| Peak twist, ° | 13.7 (11.2, 16.1) | 10.8 (8.3, 13.2) | 2.9 (0.64, 5.16) | 0.048 | 0.39 | 0.40 | 0.45 |
| Peak torsion, °/cm | 1.5 (1.2, 1.8) | 1.2 (0.96, 1.5) | 0.32 (0.056, 0.59) | 0.072 | 0.33 | 0.37 | 0.40 |
| Peak untwisting rate, °/s | -32.1 (-37.8, -26.4) | -32.8 (-38.5, -27.1) | 0.74 (-3.9, 5.4) | >0.9 | 0.49 | 0.60 | 0.36 |
| Peak twisting rate, °/s | 31.2 (25.9, 36.5) | 28.5 (23.2, 33.8) | 2.6 (-3.3, 8.7) | 0.768 | 0.21 | 0.40 | 0.32 |

Data are means (95% confidence intervals), $P_{Bon}$ are Bonferroni adjusted P values. **Abbreviations:** CS, circumferential strain; EDV, end-diastolic volume; EF, ejection fraction; ESV, end-systolic volume; GCS, global circumferential strain; GLS, global longitudinal strain; ICC, intra-class correlation coefficient ;LS, longitudinal strain; LV, left ventricular; PTS, principle tangential strain; RS, radial strain ; SV, stroke volume; 3D-STE, three-dimensional speckle tracking echocardiography. \*Averaged based on 16-segments model. <sup>†</sup>ICC represents data from good and sub-optimal images analysed together. <sup>‡</sup>ICC after restricting the analysis to undistorted quality images. <sup>§</sup>ICC after restricting the analysis to distorted quality images.

**FIGURE 2.**
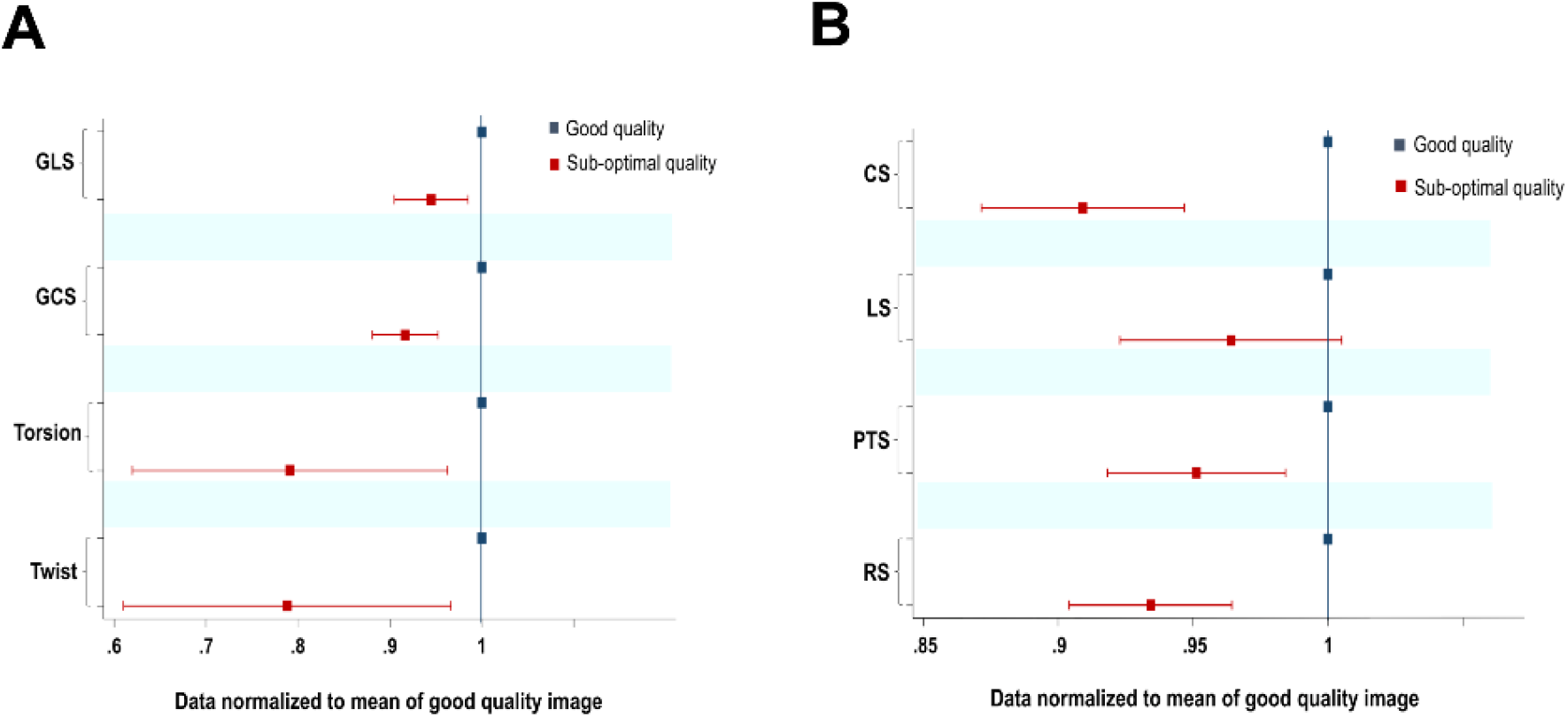
Extent of bias represented in standardized terms (Study 1). Left ventricular (LV) global deformation and rotational indices (A), and averaged segmental LV deformation indices (B) of the good and sub-optimal 3DE image quality. Data normalized to the mean of good quality image=1. CS, circumferential strain; GCS, global circumferential strain; GLS, global longitudinal strain; LS, longitudinal strain; PTS, principle tangential strain; RS, radial strain.

**FIGURE 3.**
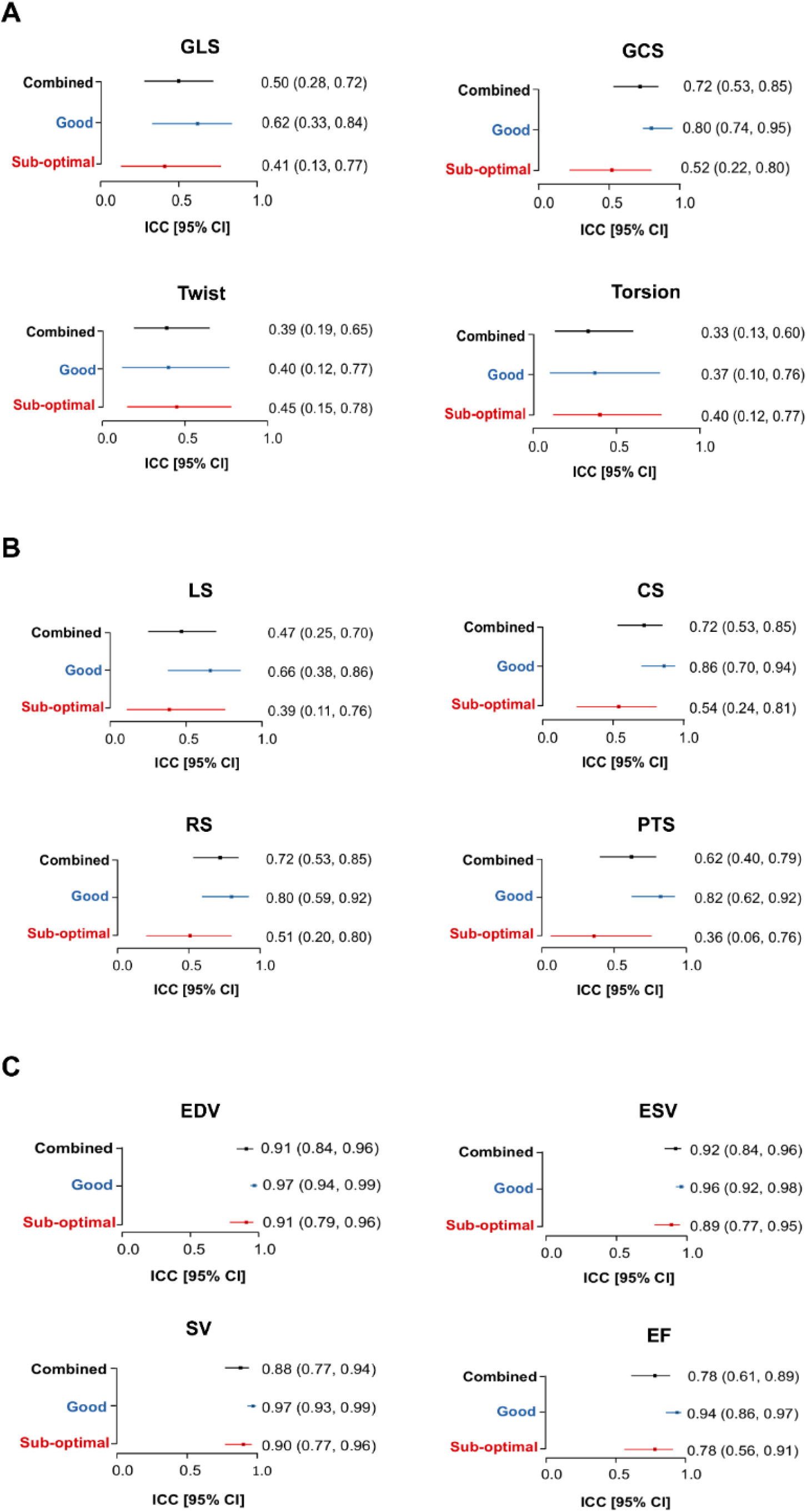
Test-retest repeatability. Intra-class correlation coefficient (ICC) of left ventricular (LV) global deformation and rotational indices (A); average segmental LV deformation indices (B); and volumetric indices (C). Combined ICC represents data from good and sup-optimal images analysed together good ICC after restricting the analysis to undistorted quality images and sub-optimal ICC after restricting the analysis to distorted quality images. Abbreviations are as in Figure 2 in addition to EDV, end-diastolic volume; EF, ejection fraction; ESV, end-systolic volume; and SV, stroke volume.

### EXTENT OF BIAS RELATIVE TO DEGREE OF DEGRADATION IN IMAGE QUALITY BY IMPAIRING ULTRASOUND PROPAGATION (STUDY 2)

Results from study 2 using the neoprene rubber confirmed the results obtained from study 1. There was a systematic bias in all global and segmental LV deformation and rotational indices. Moreover, the extent of this systematic underestimation was proportional to the extent of degradation in image quality (i.e. the poorer the image quality the larger the bias) (Table 3; Figure 4). Volumetric indices including LVEF were also progressively underestimated relative to the extent of impairment in image quality (Table 3).

**Table 3.**
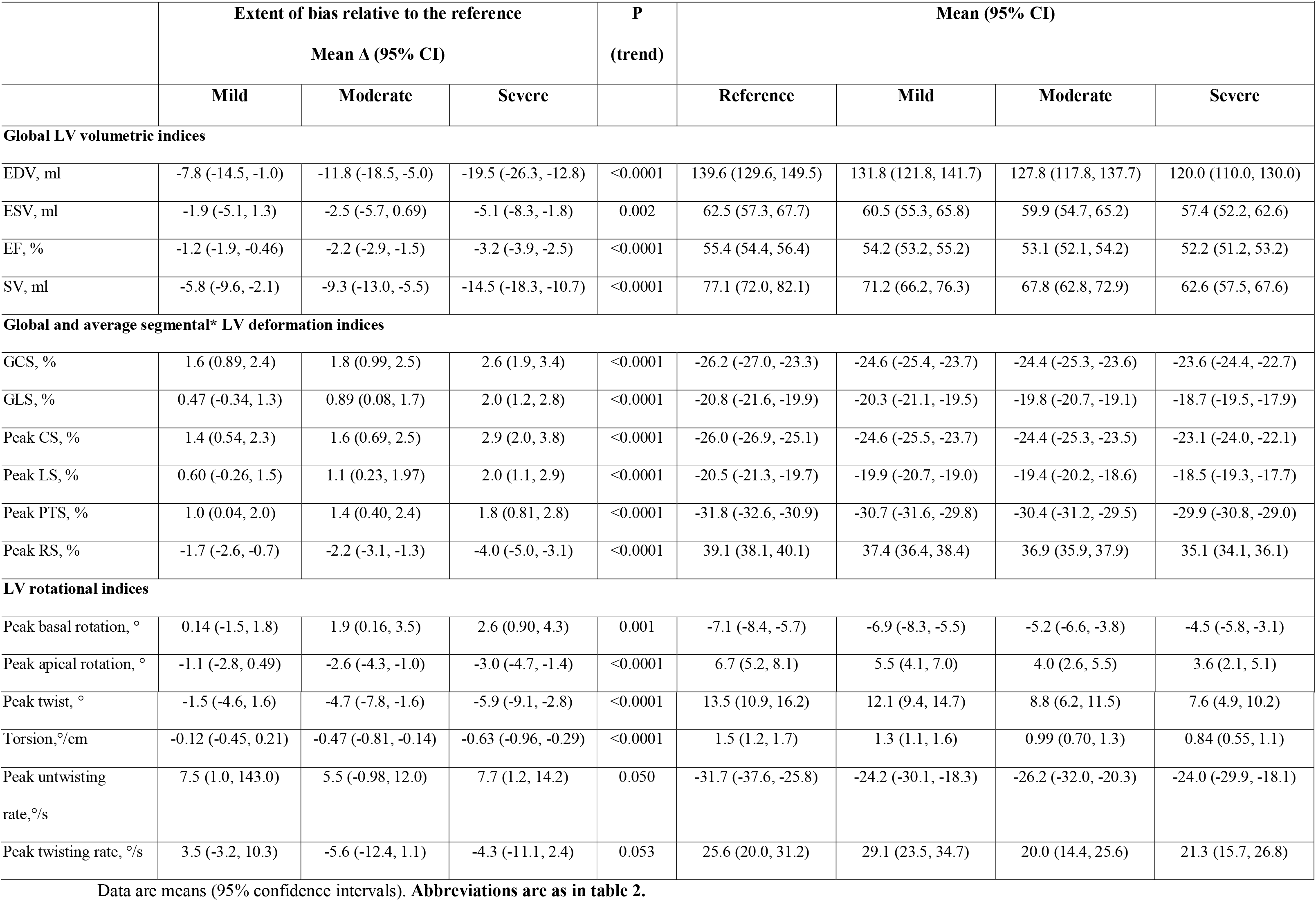
The extent of bias proportional to the impairment in image quality of 3D-STE derived LV indices (study 2)

**FIGURE 4.**
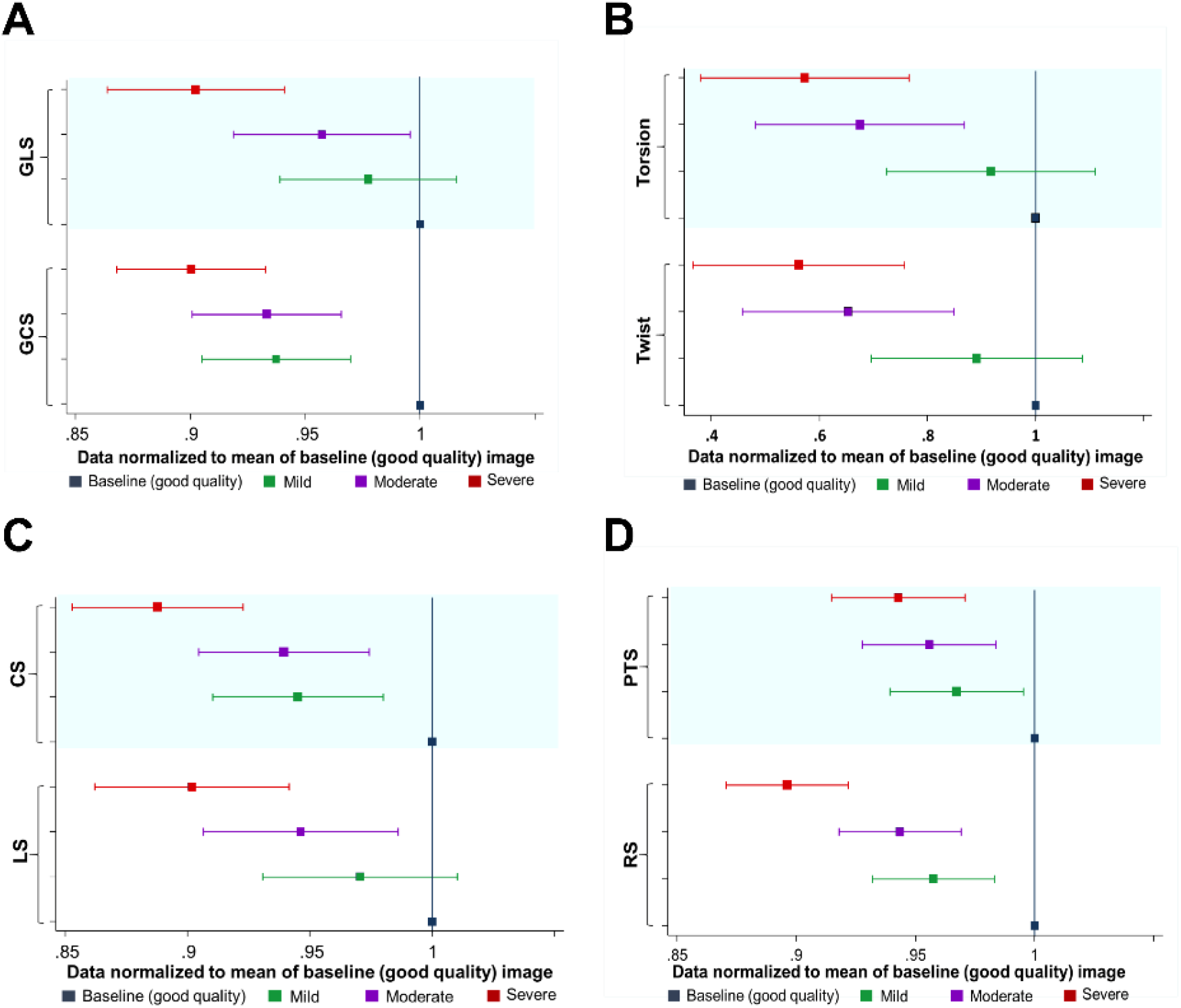
Extent of bias represented in standardized terms (Study 2). Left ventricular (LV) global deformation (A), rotational (B) and averaged segmental deformation (C, D) indices of the baseline (good quality) 3DE image and its corresponding mildly, moderately and severely impaired image quality. Data were normalized to mean of optimal image=1. Abbreviations are as in Figure 2.

## DISSCUSSION

We provide quantitative evidence regarding the impact of sub-optimal image quality on 3D-STE derived LV deformation and rotational indices. We show for the first time that sub-optimal 3DE image quality whether due to poor echocardiographic technique or to poor acoustic windows causes systematic bias (i.e. systematic underestimation of measurements). In images where quality was degraded by poor technique the magnitude of this bias was ~10% for deformation indices, ~20% for rotational indices, and <10% for volume measures. The extent of the bias was similar (or a little worse for volume indices) when an ultrasound-attenuating material was used to degrade image quality, and this study also showed that the bias was proportional to the degree of degradation in image quality (i.e. the poorer the image quality, the larger the bias). Further, image quality seems to have moderate effects on the reproducibility of 3D-STE derived indices, particularly LV deformation indices.

The impact of sub-optimal image quality on 3D-STE has previously been discussed (20–22), but no study has provided quantitative evidence on the magnitude or direction of any bias. Trache et al. (20) reported better agreement between 2D-STE and 3D-STE when sub-optimal segments were excluded (20) which is consistent with our test-retest findings. Gayat et al. (21) studied inter-vendor agreement of 3D-STE and mentioned image quality as one possible factor which might explain inter-vendor discordance; however these authors reported that image quality did not hugely affect the inter-vendor agreement based on excluding participants with sub-optimal 3DE images (21). Using subjectively scored image quality (poor, fair, good and excellent), Muraru et al. (22) reported a positive association between image quality and 3D-STE derived LV strain indices; however due to the observational nature of the study confounding by aging, disease or some other physical characteristic associated with poor image quality could not be excluded.

Using optimal images, we found a similar reliability of 3D-STE derived LV deformation, volume and rotational indices to previously published studies (21,28–30). Poor quality images had discernible albeit modest effects on the reproducibility of volumetric indices by 3D-STE but the reproducibility of strain indices was more affected by image quality. The reproducibility of rotational indices was poor to fair irrespective of image quality.

### CLINICAL IMPLICATIONS

3D-STE is increasingly used in clinical practice and it has been suggested that it may have diagnostic and prognostic value (31–34). However, since disease is commonly associated with poor image quality and poor image quality results in a systematic bias, particularly in LV deformation and rotational indices, image quality has the potential to be an important and neglected confounder in 3D-STE studies. This is particularly relevant since the size of the bias is similar to the effects reported in disease (31–34). Further, exclusion of participants with poor image quality will not obviate this bias, since selective exclusion of participants is a potential source of bias in itself.

### STUDY LIMITATIONS

This study has a number of limitations. First, we did not include another imaging method as a reference, so we cannot comment on the absolute accuracy of 3D-STE; however, the existence of bias related to image quality does not depend on the accuracy of a measurement. We did not test our approach using software from different vendors - it is possible that the extent of bias varies between different software. In future studies it will be important to examine if image quality bias varies between software from different vendors; nevertheless, TomTec software is widely used, as it is platform independent, so these findings are relevant to a wide range of published studies.

## CONCLUSION

Sub-optimal image quality introduces a substantial systematic bias (~10%) for strain indices and more for twist and torsion (~20%). The bias is proportional to the grade of impairment in 3DE image quality (i.e. the poorer the image quality the larger the bias). Sub-optimal 3DE image quality adversely affects the reproducibility of 3D-STE (on LV deformation indices in particular). Image quality has the potential to be a neglected major confounder in studies using 3D-STE.

### CLINICAL COMPETENCIES

Sub-optimal image quality introduces a substantial systematic bias in left ventricular deformation and rotational indices by 3D-STE. The bias is proportional to the grade of impairment in 3DE image quality. Bias related to image quality might have important clinical implications since its magnitude is similar to that reported in association with disease and may confound these reported associations.

### TRANSLATIONAL OUTLOOK

This study provides the first quantitative evidence regarding the impact of sub-optimal image quality on 3D-STE derived left ventricular deformation and rotational indices. Future studies are needed to examine if the reported bias varies between different vendors.

## ACKNOWLEDGEMENTS

We are grateful to all the volunteers who participated in this study.

---

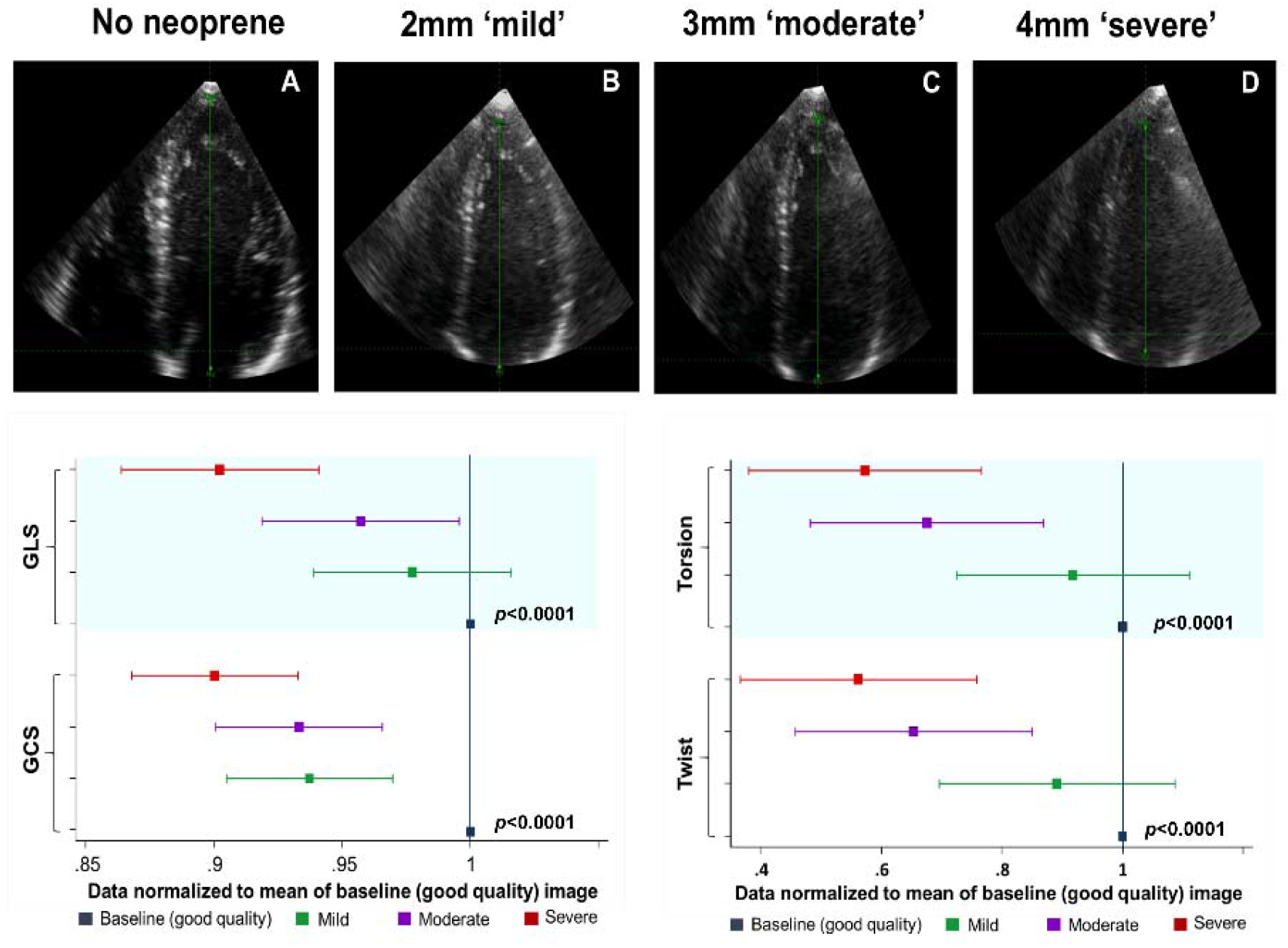
Central illustration: The impact of sub-optimal image quality on left ventricular (LV) global deformation and rotational indices by 3D spackle tracking echocardiography. An example of a 3D LV dataset with an optimal quality reference (no neoprene), mild (2mm neoprene), moderate (3mm neoprene), and severe (4mm neoprene) impairment of 3DE image quality obtained from the same participant (Top panel). The extent of the systematic bias represented in standardized terms (Bottom panel). Data normalized to mean of baseline image=1, p for trend. GLS, global longitudinal strain; GCS global circumferential strain.

## ABBREVIATIONS

CS: circumferential strain
GCS: global circumferential strain
GLS: global longitudinal strain
ICC: intraclass correlation coefficient
LS: longitudinal strain
LV: left ventricular
LVEF: left ventricular ejection fraction
PTS: principle tangential strain
RS: radial strain
STE: speckle-tracking echocardiography
3DE: three-dimensional echocardiography

